# Generative Data Augmentation for Small-Cohort Glioblastoma Diagnosis from Circulating microRNA: Why Same-Modality Validation Overstates Performance

**DOI:** 10.64898/2026.11.16.766934

**Authors:** Shivam Rane, Soheel Mishra, Phillip Carvalho, Vikas Gurjar, Rajnarayan Tiwari, Pradyumna Kumar Mishra

## Abstract

Circulating microRNAs (miRNAs) have potential as minimally invasive biomarkers for glioblastoma (GBM), but the small size of discovery groups makes it difficult to develop a classifier. The discovery cohort, GSE122488, comprised 38 patients (n = 38). Generative data augmentation was applied to increase the dataset, although evaluations based on held-out samples from the same cohort or a single validation set make it difficult to distinguish biological signals from cohort- or platform-specific effects. CTGAN and Tabular Variational Autoencoder (TVAE) models were trained on the 38-patient cohort producing 500 synthetic samples each. Random Forest and XGBoost classifiers were trained on augmented datasets and assessed using an escalating validation approach: first on an independent plasma cohort of the same modality (GSE184472, n = 118), and then on a tissue cohort from a different platform (Chinese Glioma Genome Atlas (CGGA), n = 198). XGBoost using TVAE-augmented data achieved an AUROC of 0.59 (95% CI 0.49-0.70) on GSE184472, compared with 0.42 to 0.53 without augmentation, although no AUROC on this cohort was significantly different from chance. However, this improvement did not carry to CGGA, where AUROC was 0.40 (95% CI 0.32-0.48) before harmonisation and 0.47 after Min-Max harmonisation. After harmonisation, a recall of 1.00 resulted from predicting every sample as GBM, leading to an accuracy of 0.41 and showing poor discrimination. The generalisation gap was examined in terms of modality mismatch, differing class definitions between cohorts, synthetic-data-related class imbalance, and incomplete cross-platform harmonisation. These findings show the need for independent, cross-platform validation before using data augmentation for diagnosis. The escalating validation framework provides a way to evaluate generalizability in small-cohort studies.

## 1. Introduction

Glioblastoma multiforme (GBM) is the most common and most aggressive primary malignant brain tumor in adults [1]. Circulating miRNAs in blood plasma are attractive diagnostic biomarker candidates because they are stable in circulation and obtainable via a routine blood draw rather than a surgical tissue biopsy [2]. The central obstacle to this line of research is cohort size: the discovery dataset used here, GSE122488, comprises only 38 subjects (22 tumor - 12 glioblastoma, 10 grade II-III glioma and 16 healthy controls) profiled across roughly 1,900 mature miRNAs. A classifier trained directly on 38 samples with far more features than subjects are at severe risk of learning noise rather than generalizable biology [3, 4].

Generative data augmentation synthesizing additional training examples with a learned generative model rather than collecting more patients is an increasingly common response to this problem in small-cohort biomedical machine learning [5]. Tabular generators such as Conditional Tabular Generative Adversarial Network (CTGAN) and Tabular Variational Autoencoders (TVAE) have been applied to genomic, radiomic, and other types of biological tabular data, with reported improvements in downstream classification performance [6, 7]. However, a recurring limitation across this literature concerns the evaluation method: performance is frequently assessed on a held-out portion of the same small cohort used for both training and augmentation, or on a single validation cohort obtained using the same measurement modality [6]. This leaves open a basic question: does the augmented model perform well on genuinely independent data that was collected using a different platform, or does it simply fit the idiosyncrasies of the original small cohort more accurately?

This study evaluates whether CTGAN and TVAE augmentation meaningfully improve GBM classification from circulating miRNA under an escalating, more demanding validation regime than is typical in this literature: (1) an independent cohort collected under the same modality (GSE184472, *n* = 118, blood plasma), and (2) an independent, cross-platform cohort collected under a different modality entirely (CGGA, *n* = 198, tumor tissue rather than blood, profiled on a different array platform). Evaluating on an external, cross-platform cohort is the more demanding and more clinically relevant test, and is a step that several small-cohort augmentation studies in this space skip. Our central contribution is not a new augmentation algorithm, but a transparent demonstration using standard, off-the-shelf generators and classifiers that apparent gains from small-cohort augmentation can fail to survive this more demanding evaluation, together with a concrete account of why, and what would need to change before a diagnostic claim is warranted.

Prior work on GBM molecular classification has largely relied on large reference cohorts such as TCGA to build robust classifiers from mRNA or DNA methylation data [8], or has integrated large-scale RNA-seq atlases to identify candidate regulatory networks [9]. These approaches sidestep the small-cohort problem by relying on cohort sizes in the hundreds to thousands, which are not yet available for circulating-miRNA GBM diagnostics specifically.

In the small-cohort regime, generative augmentation has been applied to glioma classification from other data modalities, including CTGAN-based augmentation of MRI radiomic features for IDH-mutation status [10] and conditional variational autoencoder augmentation of Raman spectroscopy data under a strict patient-isolated cross-validation protocol [11]. Benchmark comparisons of SMOTE, CTGAN, TVAE and diffusion-based generators on small biological tabular datasets [7], the evaluations of the usefulness of the synthetic data generally being carried out via train-on-synthetic/test-on-real protocols on data from a single source. With regard to circulating-miRNA panels for the broader purpose of glioma detection, it has been found that pooled multi-dataset panels are able to discriminate glioma cases from controls across different datasets, although it is noted that pooling leads to biological heterogeneity among glioma subtypes [12, 13].

Our work differs from this body of prior work primarily in evaluation design rather than in method: we apply standard generators (CTGAN, TVAE) and standard classifiers (Random Forest, XGBoost but evaluate the resulting pipeline against an independent, cross-platform, cross-tissue-type cohort in addition to a same-modality validation cohort. To our knowledge, this cross-platform stress test has not been reported for circulating-miRNA GBM diagnostics augmented with tabular generative models [14].

## 2. Methodology

### 2.1 Cohort and sample composition

A. *Cohorts:* Four public cohorts comprising 379 real (non-synthetic) samples were assembled **(Table 1**; **Figure 1)**: GSE122488 (n = 38), GSE184472 (n = 118), CGGA (n = 198) and GSE90603 (n = 25). Of these, 354 real samples from the first three cohorts were used in modelling. The figure of 198 refers only to the CGGA external test cohort (81 GBM and 117 non-GBM samples), not to the full study population. GSE90603 was preprocessed and intersected with the other cohorts but was not used in the modelling steps reported here; it remains available as a further external check. Synthetic samples (500 per generator, 1,000 in total) are counted separately from the real samples: they were used for model training only and were never included in any validation or test set. The figure of 379 that also appears in this manuscript in connection with TVAE refers to the number of synthetic tumour samples generated by TVAE **(Section 2.3B; Table 5)** and is unrelated to the real-sample total.
B. *Feature Harmonization:* All four cohorts were intersected on a common set of 201 mature miRNAs present across every dataset. GSE122488 and GSE184472 required GEO-header parsing corrections during preprocessing. GSE90603’s array probe IDs were translated to miRNA names via the GPL21572 annotation file, with duplicate probes collapsed by averaging. CGGA’s expression values which included some multiply-mapped probe labels (e.g., hsa-miR-199a-3p,hsa-miR-199b-3p) were parsed and aligned by sample ID against CGGA’s clinical table, with glioblastoma labels assigned from the Histology field (GBM/sGBM/rGBM = 1, all other histologies = 0).

### 2.2 Preprocessing and computational reproducibility

The data preprocessing and the development of the model were carried out separately in two Jupyter notebooks: preprocessing.ipynb and training.ipynb. The preprocessing step resulted in standardized expression matrices on a per-cohort basis and in datasets suitable for use with machine learning, limited to the 201 common mature miRNAs. The processed cohort-level matrices were saved in the data/processed/ directory, while the ML-ready discovery, validation, CGGA, and biological matrices were stored in the data/ml_ready/ directory for use in subsequent analyses. In all cases, the stochastic machine-learning and generative-model procedures employed a random_state = 42 to ensure reproducibility. The analyses were carried out on a Windows machine within a Conda-managed Python environment (environment name: torch_gpu). The main computational libraries used were scikit-learn, XGBoost, PyTorch, the standalone ctgan Python package (note: this is the independent ctgan library, not the SDV-wrapped CTGAN/TVAE implementation - relevant for anyone attempting to reproduce these results), Pandas, and NumPy. Important model parameters were set in advance and kept consistent throughout the analyses: Random Forest and XGBoost were each configured with 100 estimators/trees (maximum tree depth and learning rate were left at library defaults and not explicitly overridden); CTGAN and TVAE were each trained for 300 epochs; the Tabular Transformer baseline used a model dimension of 64, 4 attention heads, 2 encoder layers, the Adam optimizer with a learning rate of 0.001, binary cross-entropy loss, and was trained for 50 epochs on the full 38-sample training set with no mini-batching (full-batch training). The synthetic datasets were generated using 500 samples per generator (CTGAN and TVAE each), and all downstream analyses made use of the same predefined feature set and random seed (random_state = 42). Exact hardware specifications (CPU model and core count, system RAM, GPU model and VRAM) and precise software version numbers (Python, Jupyter, scikit-learn, XGBoost, PyTorch, ctgan, Pandas, NumPy, and SciPy) were not recorded at the time these notebooks were run and could not be reliably reconstructed afterward; no environment or requirements.txt was saved alongside the analysis. We note this as a limitation of the present study’s computational reporting and recommend that future work in this pipeline log the full environment (e.g. via ‘pip freez’ or a Conda environment export) at analysis time.

**Figure 1:**
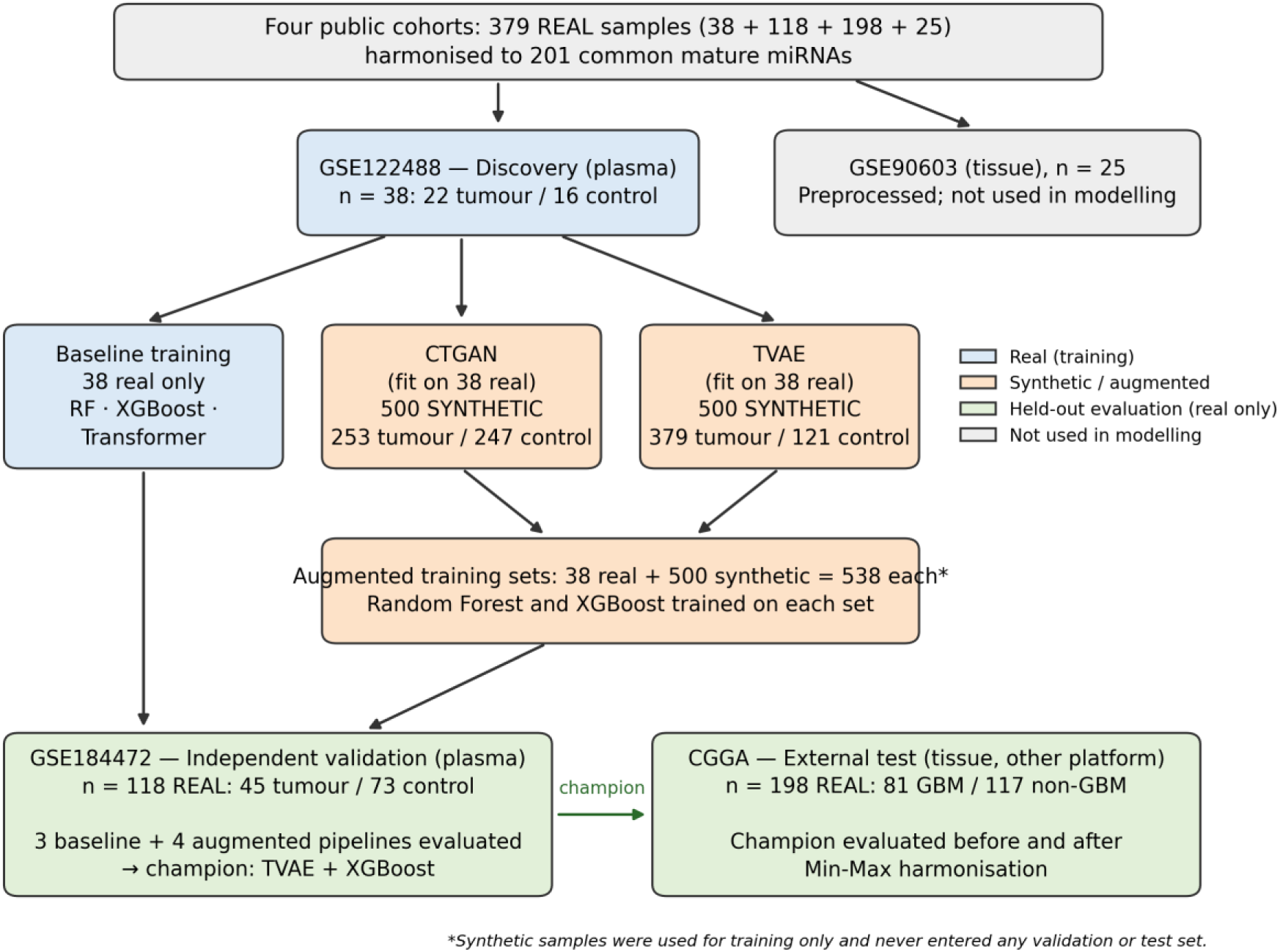
Sample flow. Real samples (37S across four cohorts) and synthetic samples (500 per generator) are shown separately. Synthetic samples were generated from, and combined with, the 38-sample discovery cohort for training only; all evaluation used real samples exclusively.

**Table 1:**
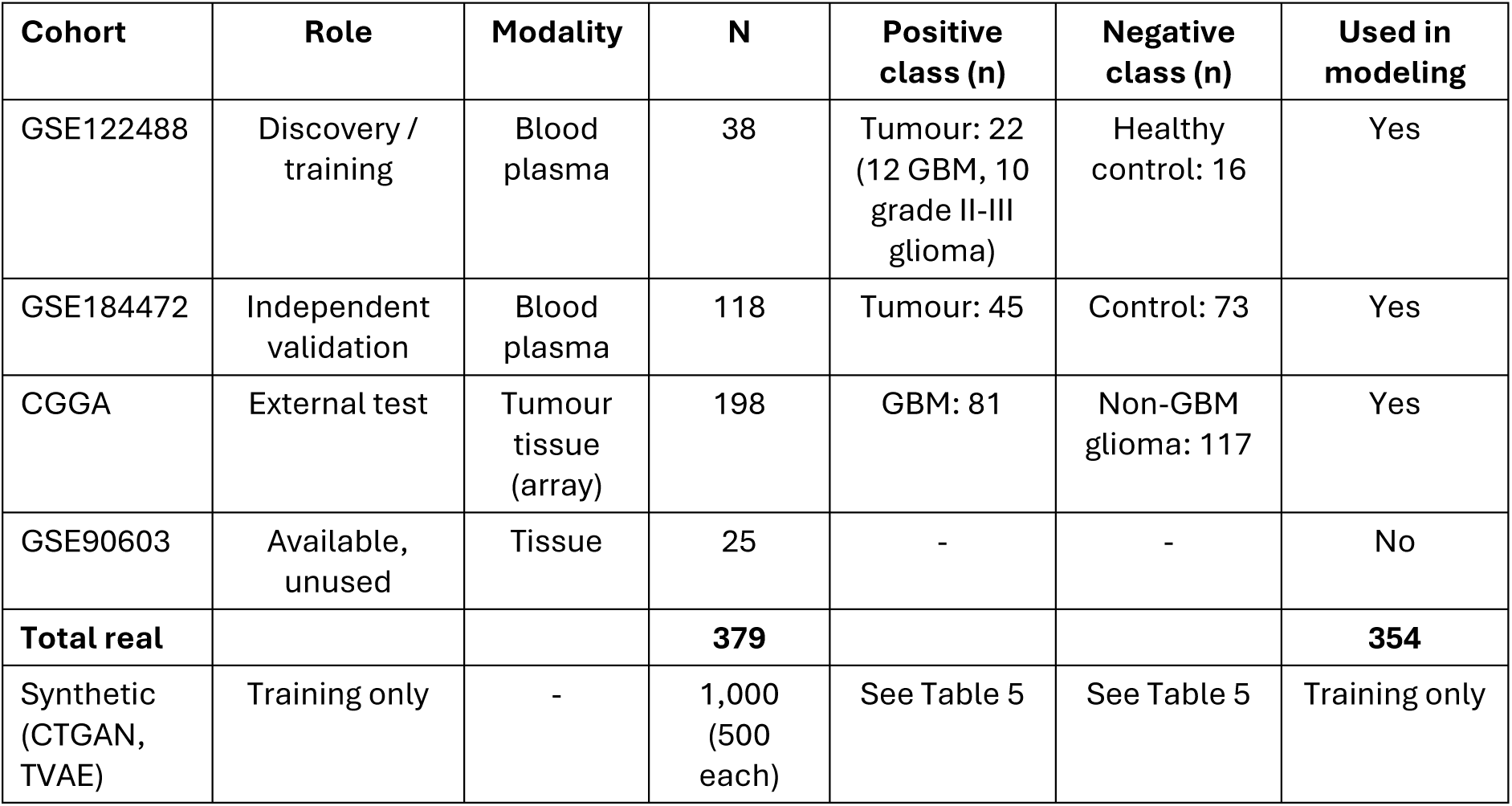
Cohorts used in this study, with class composition and use in modeling.

### 2.3 Modeling Protocol

A. *Baseline Models (No Augmentation):* Baseline performance was first established by training Random Forest, XGBoost and a PyTorch tabular Transformer solely on the 38-subject discovery cohort and evaluating them on the independent GSE184472 validation set, with no synthetic data added.
B. *Generative Augmentation:* CTGAN and TVAE were each fit on the 38-subject discovery cohort and used to generate 500 synthetic samples. CTGAN produced a near-balanced synthetic class split (253 tumor / 247 control). TVAE produced a more imbalanced split (379 tumor / 121 control). We discuss this difference in Sections 3.4 and 4 as a plausible contributor to the champion model’s downstream behavior on CGGA. Random Forest and XGBoost were retrained on each augmented set and evaluated on GSE184472 [14]. In both cases, the synthetic samples were combined with the 38 real discovery samples to form an augmented training set of 538 samples **(Figure 1)**; synthetic samples were never used for validation or testing.
C. *External Validation and Harmonization:* The champion model (best-performing model) from the augmentation experiments (TVAE combined with XGBoost) was applied to the CGGA cohort. A preliminary check of the feature order confirmed that the 201 miRNA columns were exactly matched between the training and CGGA matrices, excluding any column-alignment bug. GSE122488 (discovery) contains 22 tumor samples (12 glioblastoma, 10 grade II-III glioma) and 16 healthy controls. GSE184472 (independent validation) contains 45 tumors and 73 control samples, collected under the same blood-plasma modality as the discovery cohort. CGGA (external test) contains 81 GBM and 117 non-GBM samples of tumour tissue profiled on a different array platform; its negative class comprises all other glioma histologies rather than healthy tissue. Because the feature-order check excluded column misalignment, the performance gap was instead traced to a scale mismatch between the two data sources (training data range 0-47,580 vs. CGGA range *−*129.1-28,912.2). A Min-Max scaling step, fit independently on each cohort, was applied to both the training data (before retraining the champion model) and the CGGA cohort, and the harmonized model was re-evaluated.
D. *Biomarker Discovery:* Feature importances (Gini) were extracted from the champion model (TVAE + XGBoost, preharmonization) to identify candidate diagnostic miRNAs.

### 2.4 Statistical Evaluation

All metrics were computed on real samples only, at the default probability threshold of 0.5. From the confusion matrix of each pipeline (true positives, TP; false negatives, FN; false positives, FP; true negatives, TN) we report sensitivity (recall; TP/(TP+FN)), specificity (TN/(TN+FP)), precision (positive predictive value; TP/(TP+FP)) and F1-score (2TP/(2TP+FP+FN)), with 95% confidence intervals (CIs) for proportions calculated using the Wilson score method [15]. Because both external cohorts are class-imbalanced (tumour prevalence 0.381 in GSE184472; GBM prevalence 0.409 in CGGA), accuracy was compared with the no-information rate (NIR), i.e., the accuracy obtained by always predicting the majority class (0.619 and 0.591, respectively), using a one-sided exact binomial test.

For AUROC, 95% CIs were estimated from the Hanley-McNeil standard error, and each AUROC was compared with chance (0.5) using a two-sided z-test [16]. AUROC 95% CIs were additionally estimated by stratified bootstrap (2,000 resamples), and paired AUROC comparisons (each augmented pipeline versus its non-augmented counterpart on the same GSE184472 samples, and CGGA before versus after harmonisation) were performed with DeLong’s test for correlated ROC curves [17]. Because the exact per-sample prediction scores from the original training run were not retained (Section 2.2), the bootstrap and DeLong statistics were computed on a documented reproduction of the pipeline (reviewer_figures.py, identical real data, hyperparameters and random seeds) and are reported alongside the Hanley-McNeil-based results above as a robustness check rather than a replacement for them. Precision-recall (PR) curves were summarised by average precision (AP) and plotted against the prevalence baseline, the expected precision of a non-informative classifier, which is the appropriate reference for imbalanced data [18]. Discrimination was further examined by plotting predicted tumour probabilities separately for each true class.

Class ratios of the synthetic datasets were compared with that of the real discovery cohort using exact binomial tests. Real and synthetic miRNA distributions were compared feature-wise using two-sample Kolmogorov-Smirnov tests with Benjamini-Hochberg false-discovery-rate correction [19], by projecting real and synthetic samples onto the first two principal components of the real data, and by comparing pairwise miRNA correlation structure.

## 3. Results

### 3.1. Baseline Classification and External Generalizability

The first analysis looked into whether conventional machine-learning models, which had been trained directly on the 38-subject GSE122488 discovery cohort, would be able to generalize to an independent cohort that had been collected using the same biological modality. For this purpose, Random Forest, XGBoost, and a PyTorch tabular Transformer were trained without the use of synthetic augmentation and tested on GSE184472, an independent plasma cohort consisting of 118 samples. All three of the baseline models showed only limited discriminative ability **(Table 2).** The Random Forest had the highest baseline Area Under the Receiver Operating Characteristic Curve (AUROC) at 0.527, together with an accuracy of 0.415 and a recall of 0.978. XGBoost obtained an AUROC of 0.459, an accuracy of 0.407, and a recall of 0.689, whereas the Transformer achieved an AUROC of 0.421, an accuracy of 0.415, and a recall of 0.667. Thus, despite the relatively high recall obtained by Random Forest, its AUROC remained close to 0.50 and its accuracy was low. The fact that there was a discrepancy between recall and AUROC shows that the high sensitivity was not linked to reliable separation of the classes. In general, the baseline models offered little evidence to suggest that the 38-subject discovery cohort on its own contained enough information for robust out-of-cohort classification.

**Table 2:** Baseline performance of non-augmented classifiers on the independent same-modality GSE184472 cohort.

| **Model** | **Accuracy** | **Recall** | **AUROC (95% CI)** |
|---|---|---|---|
| Random Forest | 0.415 | 0.978 | 0.527 (0.419-0.635) |
| XGBoost | 0.407 | 0.689 | 0.459 (0.352-0.566) |
| Transformer | 0.415 | 0.667 | 0.421 (0.316-0.526) |

### 3.2. Generative Augmentation and Model Performance

In order to find out whether synthetic data could enhance generalisation based on the small discovery cohort, CTGAN and TVAE were each trained using GSE122488 and produced 500 synthetic observations as a result. The resulting datasets were then used to train Random Forest and XGBoost classifiers, after which the classifiers were evaluated on the independent GSE184472 plasma cohort. The four augmented pipelines showed quite different performance profiles **(Table 3).** CTGAN combined with Random Forest reached an accuracy of 0.559, a recall of 0.578, and an AUROC of 0.573. CTGAN together with XGBoost obtained the highest accuracy of all the augmented models (0.661), but this was associated with a very low recall of 0.133 and an AUROC of only 0.553. Hence, the seemingly positive accuracy of this model did not result in the adequate identification of tumor samples. The TVAE combined with Random Forest resulted in a recall of 1.000 but had an accuracy of just 0.381 and an AUROC of 0.528, demonstrating that complete recovery of the positive class can occur without meaningful discrimination. In contrast, TVAE + XGBoost obtained the highest AUROC of all the four augmented pipelines, attaining a value of 0.592, with an accuracy of 0.542 and a recall of 0.622. The TVAE + XGBoost model thus proved to be the best augmented configuration within the predefined same-modality validation framework and was therefore named the champion model for the subsequent external testing. Yet its AUROC value of 0.592 indicates only modest discriminatory ability and must not be seen as proof of clinically useful diagnostic performance.

**Table 3:** Performance of generatively augmented classifiers on independent same-modality GSE184472 validation.

| **Augmentation-classifier pipeline** | **Accuracy** | **Recall** | **AUROC (95% CI)** |
|---|---|---|---|
| CTGAN + Random Forest | 0.559 | 0.578 | 0.573 (0.465-0.681) |
| CTGAN + XGBoost | 0.661 | 0.133 | 0.553 (0.445-0.661) |
| TVAE + Random Forest | 0.381 | 1.000 | 0.528 (0.420-0.636) |
| **TVAE + XGBoost** | **0.542** | **0.622** | **0.592 (0.485-0.699)** |

Importantly, comparison with the non-augmented XGBoost baseline showed an increase in AUROC from 0.459 to 0.592 following TVAE augmentation. This represents an absolute improvement of 0.133 AUROC units on the independent plasma cohort. Yet since the obtained AUROC was still considerably lower than what would be expected for a reliable diagnostic classifier, the result was seen as indicating only a modest gain in same-modality performance rather than as proof of genuine diagnostic improvement.

### 3.3. Confusion-Matrix Metrics, Uncertainty and Threshold-Independent Performance

To characterise performance beyond accuracy, recall and AUROC, confusion-matrix-derived metrics with 95% CIs were computed for every pipeline on GSE184472 and for the champion model on CGGA **(Table 4**; **Figure 4**; **Figure 8)**. Three observations follow. First, no pipeline achieved accuracy significantly above the NIR of 0.619 on GSE184472. The only pipeline whose accuracy exceeded the NIR, CTGAN + XGBoost (0.661; p = 0.197), did so almost entirely by predicting the control class (specificity 0.986, sensitivity 0.133; 7 of 118 samples predicted positive). Second, two pipelines collapsed towards uniform positive prediction: the Random Forest baseline (specificity 0.068) and TVAE + Random Forest, which classified every sample as tumour (specificity 0.000). Their high recall therefore carries no discriminative information, and their precision equals or approaches the prevalence (0.381). Third, TVAE + XGBoost provided the most balanced operating point: sensitivity 0.622 (95% CI 0.476-0.749), specificity 0.493 (0.382-0.605), precision 0.431 (0.318-0.552) against a prevalence of 0.381, and F1-score 0.509.

**Table 4:** Confusion-matrix metrics with S5% CIs for all pipelines on GSE184472 and for the champion model on CGGA. Threshold = 0.5. Proportion CIs: Wilson score; AUROC CIs: Hanley-McNeil. NIR, no-information rate (accuracy of always predicting the majority class); p, one-sided exact binomial test of accuracy > NIR. CGGA negative class = non-GBM glioma tissue.

| Pipeline | TP/FN/FP/TN | Sensitivity (95% CI) | Specificity (95% CI) | Precision (95% CI) | F1 | AUROC (95% CI) | Accuracy (p vs NIR) |
|---|---|---|---|---|---|---|---|
| **GSE184472 (n = 118; 45 tumour / 73 control; prevalence 0.381; NIR 0.619)** |  |  |  |  |  |  |  |
| Random Forest (baseline) | 44/1/68/5 | 0.978 (0.884-0.996) | 0.068 (0.030-0.151) | 0.393 (0.307-0.485) | 0.561 | 0.527 (0.419-0.635) | 0.415 (p = 1.000) |
| XGBoost (baseline) | 31/14/56/17 | 0.689 (0.543-0.805) | 0.233 (0.151-0.342) | 0.356 (0.264-0.461) | 0.470 | 0.459 (0.352-0.566) | 0.407 (p = 1.000) |
| Transformer (baseline) | 30/15/54/19 | 0.667 (0.521-0.786) | 0.260 (0.173-0.371) | 0.357 (0.263-0.464) | 0.465 | 0.421 (0.316-0.526) | 0.415 (p = 1.000) |
| CTGAN + Random Forest | 26/19/33/40 | 0.578 (0.433-0.710) | 0.548 (0.434-0.657) | 0.441 (0.322-0.567) | 0.500 | 0.573 (0.465-0.681) | 0.559 (p = 0.921) |
| CTGAN + XGBoost | 6/39/1/72 | 0.133 (0.063-0.262) | 0.986 (0.926-0.998) | 0.857 (0.487-0.974) | 0.231 | 0.553 (0.445-0.661) | 0.661 (p = 0.197) |
| TVAE + Random Forest | 45/0/73/0 | 1.000 (0.921-1.000) | 0.000 (0.000-0.050) | 0.381 (0.299-0.471) | 0.552 | 0.528 (0.420-0.636) | 0.381 (p = 1.000) |
| TVAE + XGBoost | 28/17/37/36 | 0.622 (0.476-0.749) | 0.493 (0.382-0.605) | 0.431 (0.318-0.552) | 0.509 | 0.592 (0.485-0.699) | 0.542 (p = 0.963) |
| **CGGA (n = 198; 81 GBM / 117 non-GBM; prevalence 0.409; NIR 0.591)** |  |  |  |  |  |  |  |
| TVAE + XGBoost, before harmonisation | 6/75/13/104 | 0.074 (0.034-0.152) | 0.889 (0.819-0.934) | 0.316 (0.154-0.540) | 0.120 | 0.403 (0.324-0.482) | 0.556 (p = 0.861) |
| TVAE + XGBoost, after Min-Max harmonisation | 81/0/117/0 | 1.000 (0.955-1.000) | 0.000 (0.000-0.032) | 0.409 (0.343-0.479) | 0.581 | 0.467 (0.386-0.548) | 0.409 (p = 1.000) |

Every AUROC 95% CI on GSE184472 included 0.5 **(Table 4)**. The champion model’s AUROC of 0.592 (95% CI 0.485-0.699) did not differ significantly from chance (p = 0.092). A conservative comparison with the non-augmented XGBoost baseline (AUROC 0.459, 95% CI 0.352-0.566), which ignores the positive correlation between ROC curves estimated on the same samples, gave an AUROC difference of 0.133 (z = 1.73, p = 0.084). The gain from TVAE augmentation is therefore suggestive but not statistically established at this sample size. On the reproduction run, bootstrap AUROC CIs agreed closely with the Hanley-McNeil intervals reported above (within ±0.02 AUROC units for every pipeline). DeLong tests for the four augmented-versus-baseline comparisons (CTGAN + Random Forest vs. Random Forest, CTGAN + XGBoost vs. XGBoost, TVAE + Random Forest vs. Random Forest, and TVAE + XGBoost vs. XGBoost) gave p-values ranging from 0.03 to 0.10, and the comparison of CGGA AUROC before versus after Min-Max harmonisation gave p = 0.72. These values are in the same direction as, and of comparable magnitude to, the conservative comparison above; given that four comparisons were made and that the result depends on a reproduction rather than the original per-sample scores, we do not treat any individual comparison as establishing a robust augmentation effect. ROC and PR curves for all pipelines are shown in **Figures 2 and 3**, and the distributions of predicted probabilities by true class in **Figure 5**. As **Figure 2** shows, every ROC curve for GSE184472 (panel A) repeatedly crosses the diagonal rather than lying consistently above it, consistent with the AUROCs close to 0.5 reported in **Tables 2-4**; none shows the sustained upper-left bow expected of a strong classifier. On CGGA (panel B), the champion model’s curve also remains close to chance both before and after Min-Max harmonisation. In the precision-recall curves **(Figure 3)**, average precision exceeds the prevalence baseline for most pipelines on GSE184472, but only modestly, and for the champion model on CGGA average precision stays close to the 0.409 baseline in both settings. The predicted-probability distributions **(Figure 5)** show substantial overlap between the tumour and control classes for every pipeline, with no pipeline showing the clearly bimodal, well-separated distributions that would indicate reliable discrimination; after CGGA harmonisation, both classes’ predicted probabilities concentrate near 1.0, consistent with the universal-positive-prediction pattern described below.

**Figure 2:**
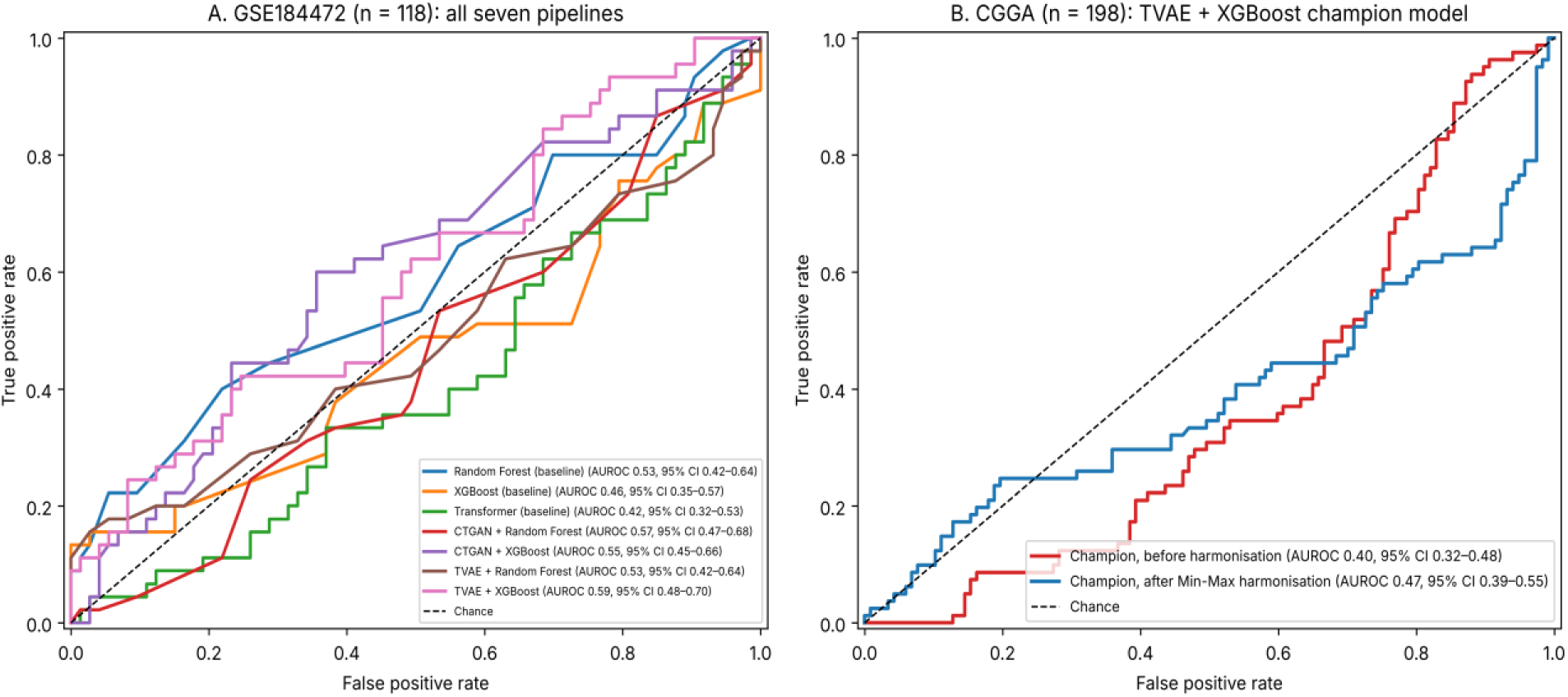
Receiver operating characteristic (ROC) curves for (A) the three baseline and four augmented pipelines on GSE184472 and (B) the TVAE + XGBoost champion model on CGGA before and after Min-Max harmonisation. AUROC values with S5% CIs are given in the legend; the diagonal indicates chance performance.

**Figure 3:**
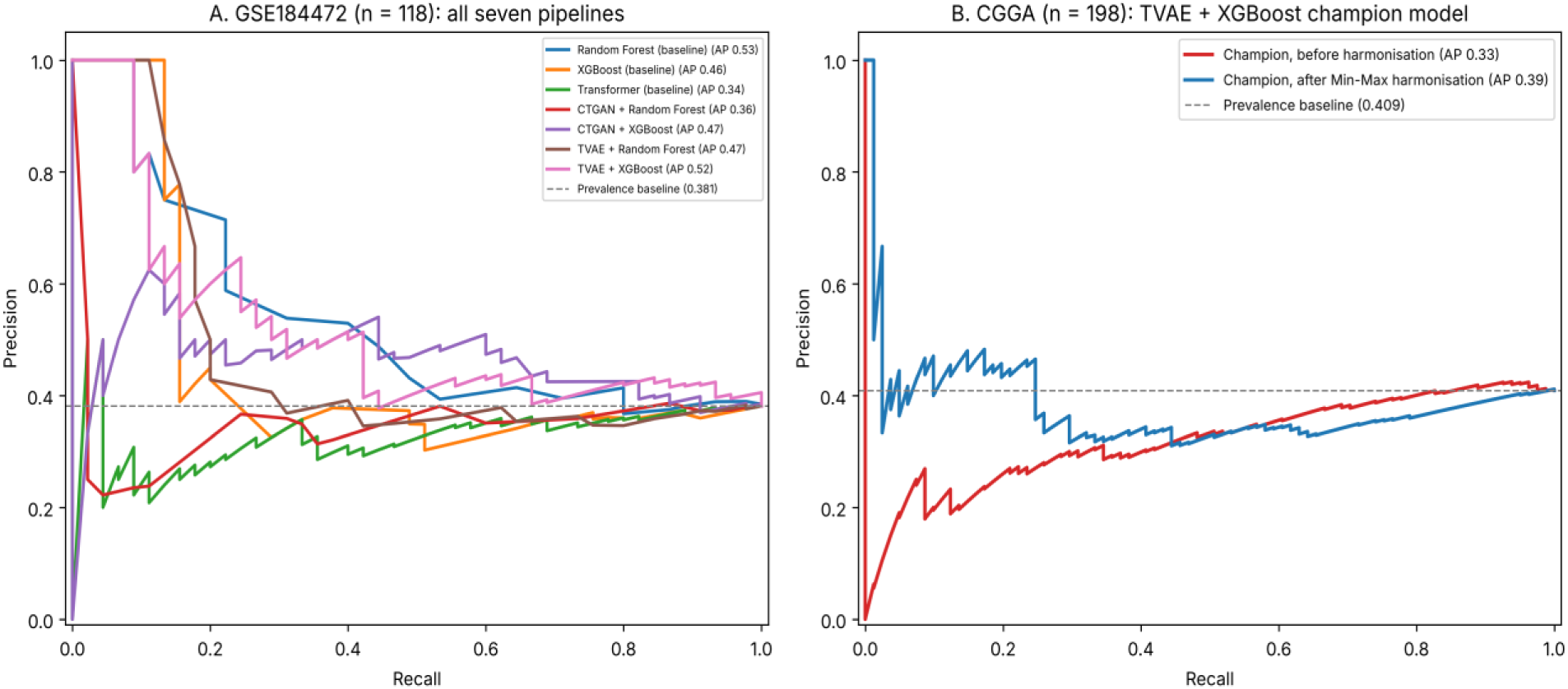
Precision-recall curves for the same models and cohorts as Figure 2. Dashed horizontal lines mark the prevalence baseline (0.381 for GSE184472; 0.40S for CGGA), the expected precision of a non-informative classifier; average precision (AP) is given in the legend.

**Figure 4:**
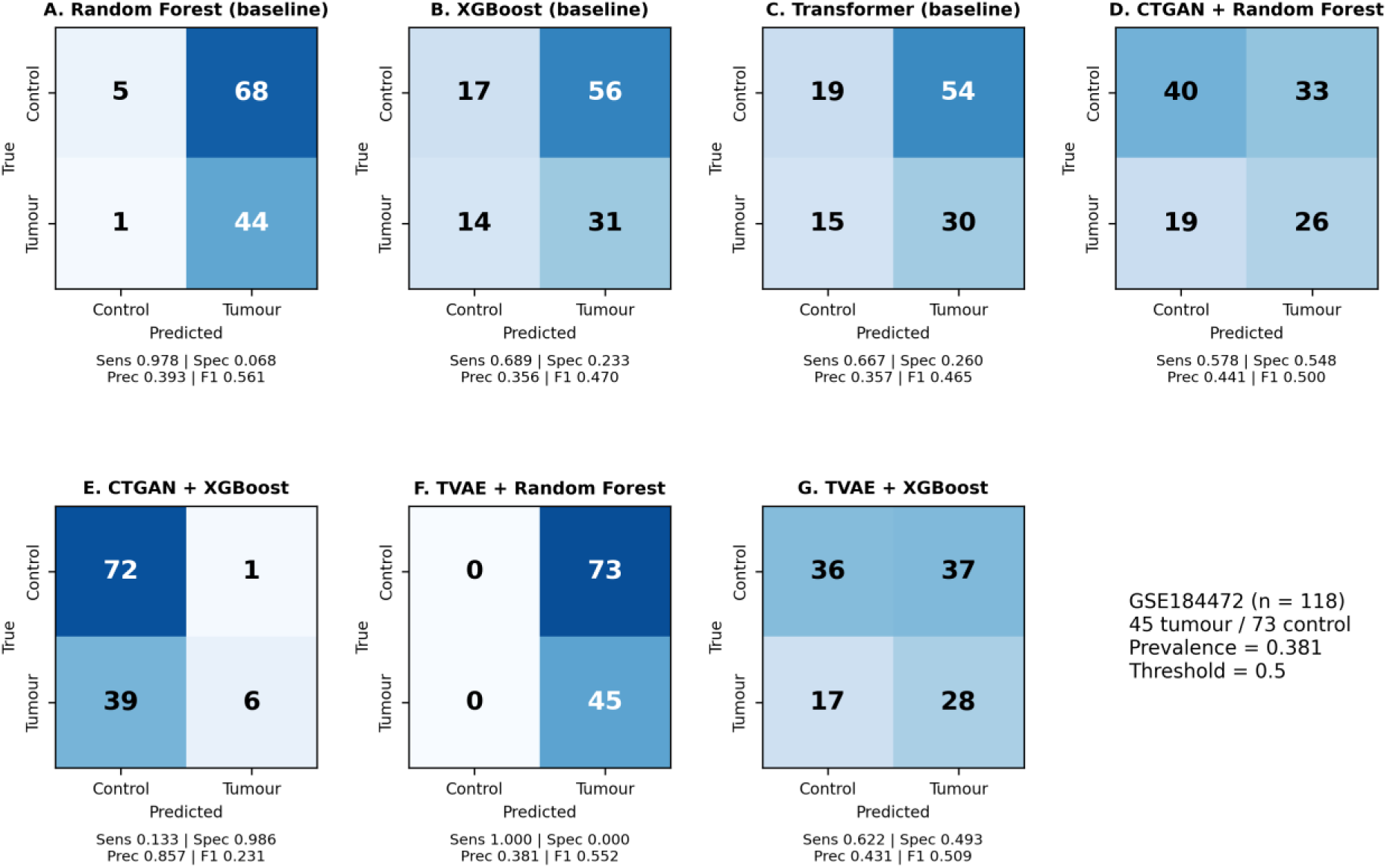
Confusion matrices of all seven pipelines on the independent GSE184472 plasma cohort (n = 118; 45 tumour, 73 control) at a 0.5 threshold, annotated with sensitivity (Sens), specificity (Spec), precision (Prec) and F1-score. Panels A-C, baseline models; D-G, augmented pipelines.

**Figure 5:**
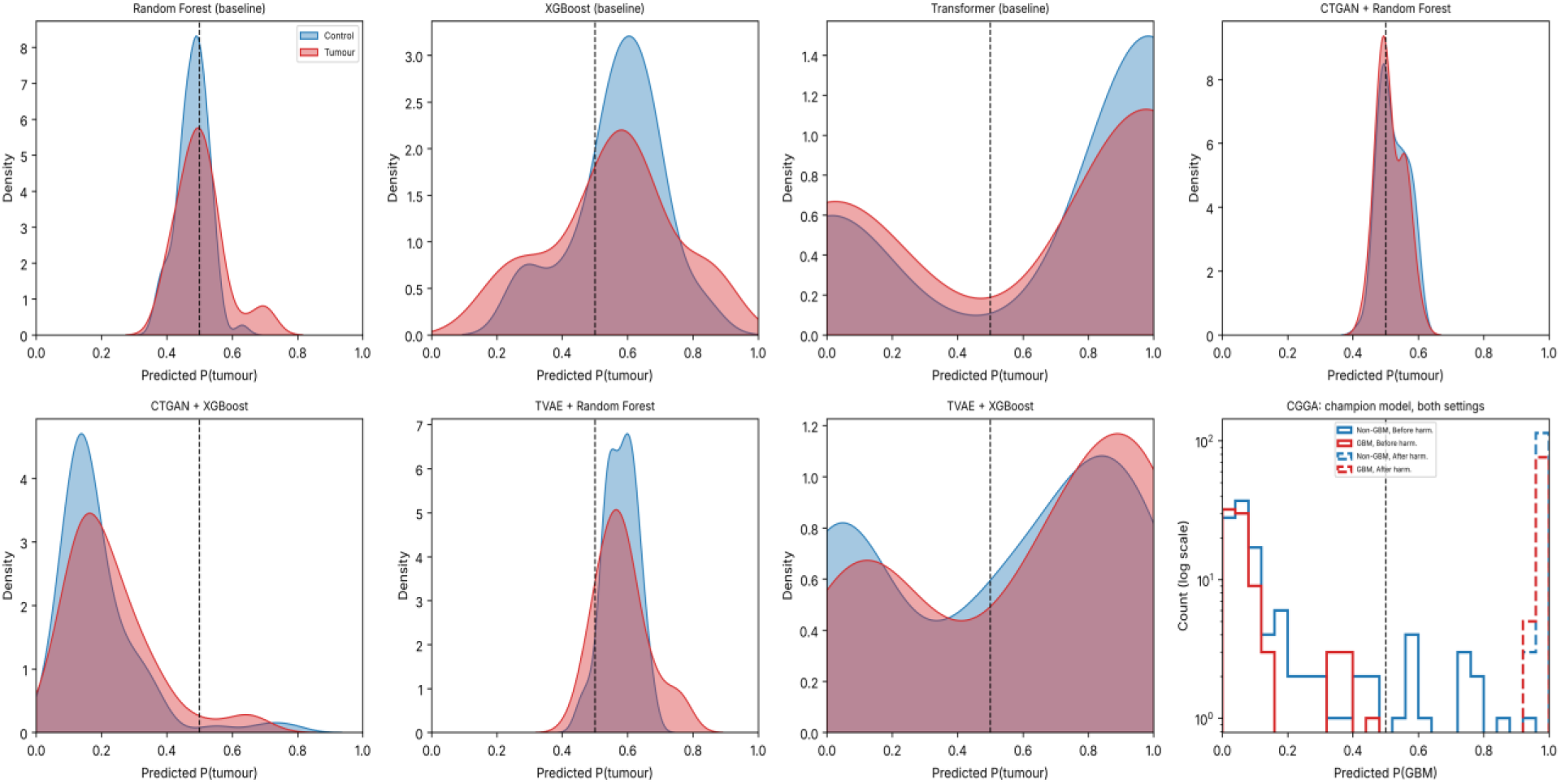
Predicted tumour probability by true class for (A) the seven pipelines on GSE184472 and (B) the champion model on CGGA before and after harmonisation. The dashed line marks the 0.5 decision threshold; well-separated distributions indicate good tumour-control discrimination.

### 3.4. Synthetic Cohort Structure, Class Distribution and Fidelity

Even though both generative models produced 500 synthetic observations, CTGAN generated 253 tumor cases and 247 control samples, corresponding to an approximately balanced synthetic dataset. In contrast, TVAE generated 379 tumor and 121 control samples, resulting in a pronounced tumor-to-control imbalance. This difference is relevant to interpretation of downstream classifier behavior because the TVAE-generated training distribution contained approximately three times as many synthetic tumor samples as control samples. Even though the current analysis does not prove a causal relationship between synthetic class composition and model decision behavior, the difference represents an important distributional characteristic of the augmentation procedure and offers a reasonable explanation for the changed positive-class behaviour seen in the later external testing. Hence, the fact that TVAE + XGBoost performs better on GSE184472 cannot be explained entirely by the generative quality of TVAE; it also has to be understood in the context of the considerably different synthetic class distribution produced by the generator.

**Table 5** summarizes these class ratios. Relative to the real discovery cohort (57.9% tumour), CTGAN shifted the class ratio towards balance (50.6% tumour; exact binomial p = 0.001), whereas TVAE over-represented the tumour class (75.8%; p < 0.001). Consequently, the TVAE-augmented training set was 74.5% tumour, compared with 51.1% for CTGAN. For CTGAN, 189 of the 201 common miRNAs (94.0%) showed distributions significantly different from the real data after Benjamini-Hochberg FDR correction; for TVAE, 160 of 201 (79.6%) did so **(Figure 6A-C)**. In a PCA space fitted on the real discovery cohort, 79% of CTGAN-synthetic samples and 99% of TVAE-synthetic samples fell within the real data’s 95% confidence region **(Figure 6A)**, indicating that both generators reproduced the real cohort’s broad low-dimensional structure despite the high rate of feature-wise distributional difference. Agreement in pairwise miRNA correlation structure (Pearson correlation between real and synthetic feature-correlation matrices) was weak for CTGAN (r = -0.01) and moderate for TVAE (r = 0.45), indicating that TVAE preserved more of the real data’s correlation structure than CTGAN did; this is consistent with TVAE’s comparatively stronger downstream performance in Sections 3.2-3.3.

**Table 5:** Class composition of real, synthetic and augmented training datasets. *Exact binomial test of the synthetic tumour proportion against the real discovery-cohort proportion (22/38).

| **Dataset** | **n** | **Tumour** | **Control** | **% tumour** | **Tumour : control** | **p vs real ratio\*** |
|---|---|---|---|---|---|---|
| Real discovery cohort (GSE122488) | 38 | 22 | 16 | 57.9% | 1.38 : 1 | - |
| CTGAN synthetic | 500 | 253 | 247 | 50.6% | 1.02 : 1 | 0.001 |
| TVAE synthetic | 500 | 379 | 121 | 75.8% | 3.13 : 1 | < 0.001 |
| CTGAN augmented training set (real + synthetic) | 538 | 275 | 263 | 51.1% | 1.05 : 1 | - |
| TVAE augmented training set (real + synthetic) | 538 | 401 | 137 | 74.5% | 2.93 : 1 | - |

**Figure 6:**
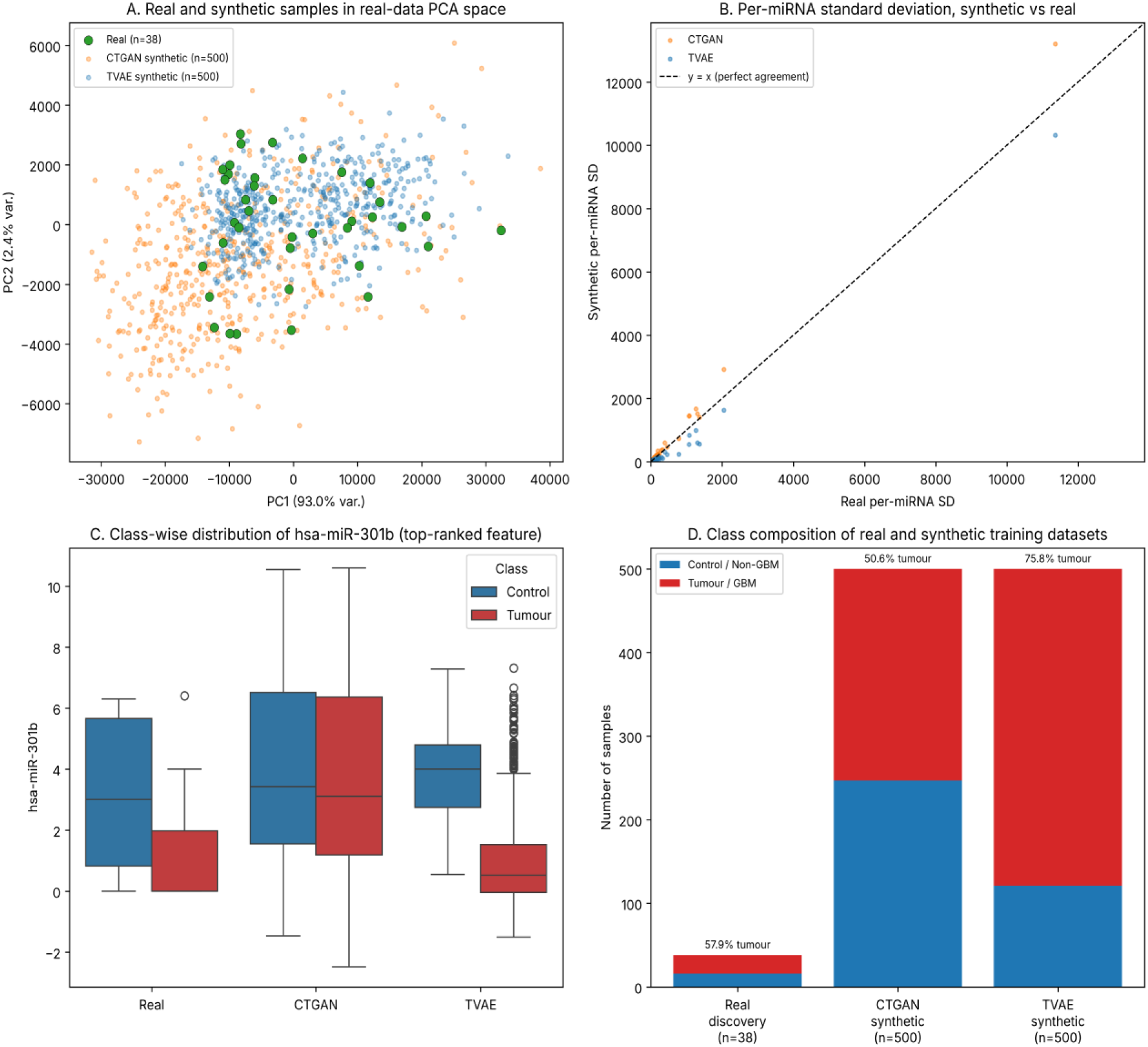
Comparison of real and synthetic miRNA distributions. (A) Real (GSE122488, n = 38) and synthetic (CTGAN, TVAE; n = 500 each) samples projected onto the first two principal components of the real data; (B) per-miRNA mean and standard deviation of synthetic versus real data; (C) class-wise distributions of the top-ranked miRNAs; (D) class composition of each dataset.

### 3.5. Model-Derived miRNA Feature Signatures

In the TVAE + XGBoost champion model, an examination of feature importance was carried out in order to find the miRNAs that contributed most strongly to classification **(Figure 7)**. The model showed a highly non-uniform distribution of importance among the 201 common miRNA features. hsa-miR-301b was the most important feature, its Gini importance being 0.183. The feature that came in second place was hsa-miR-760, with an importance value of 0.049. Hence, hsa-miR-301b had more than 3.5 times the importance of the second-ranked feature. It is noteworthy that one feature predominates so greatly despite the small size of the original discovery group and the subsequent use of synthetic augmentation. Yet the fact that a single model has identified a particular feature does not prove that it has biological or diagnostic validity. The hsa-miR-301b signal was therefore regarded as a candidate feature derived from a model and thus required further independent verification, not considered a validated biomarker.

**Figure 7:**
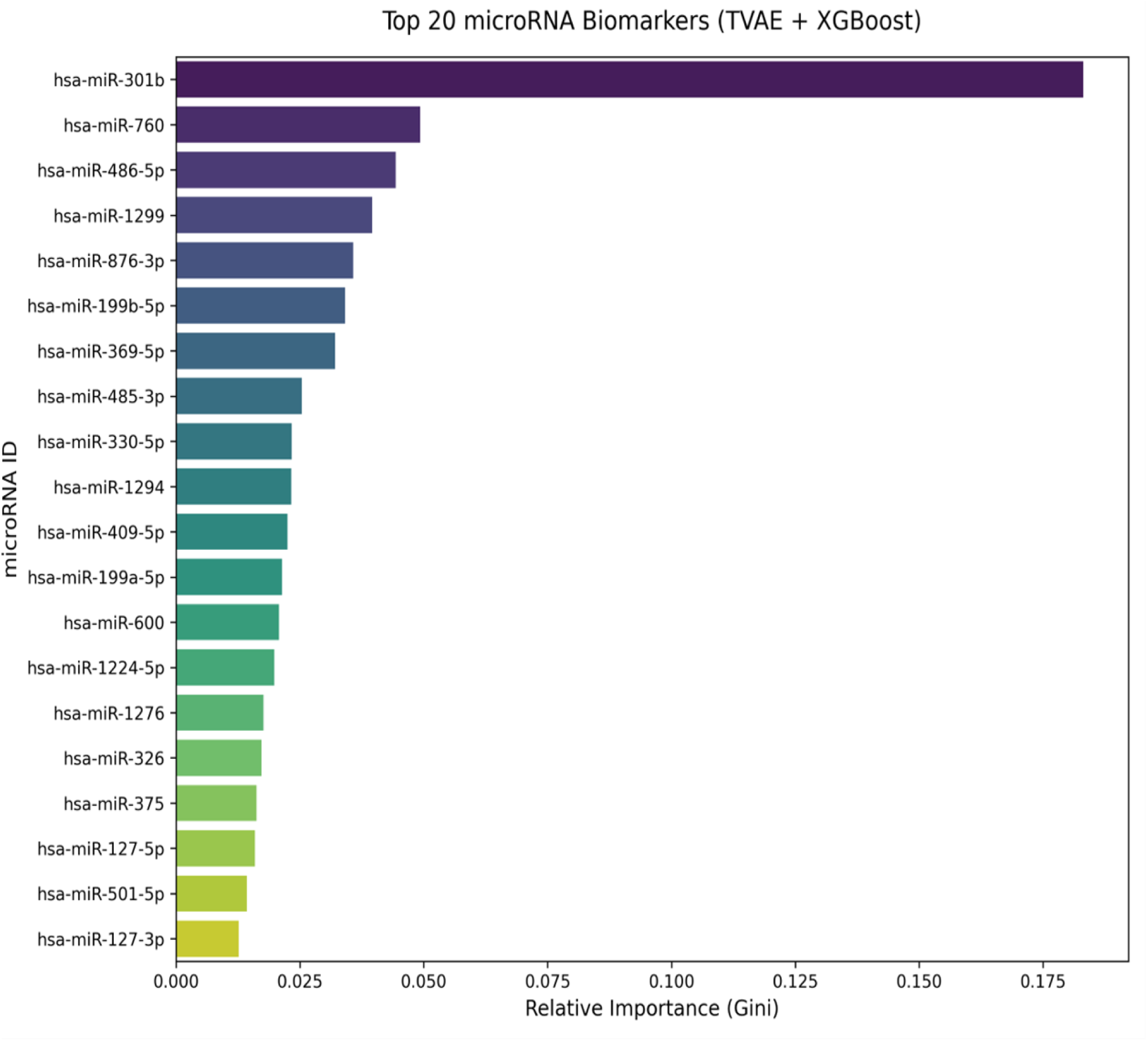
Top 20 miRNAs ranked according to Gini feature importance in the TVAE + XGBoost champion model.

### 3.6. Cross-Domain Evaluation of the Champion Model

The champion model’s robustness was then looked at in a more rigorous external validation situation involving the CGGA cohort (n = 198). Different from the discovery and GSE184472 cohorts, which included measurements of circulating plasma miRNA, the CGGA cohort consisted of tumor-tissue miRNA data obtained by means of a different measurement platform. This assessment thus involved both a biological compartment shift and a platform shift.

Before harmonization, direct application of the TVAE + XGBoost model to CGGA resulted in a substantial deterioration in performance **(Table 6**; **Figure 8A)**. Accuracy was 0.556, the recall dropped sharply to 0.074 and the AUROC was 0.403. The low recall shows that only a small proportion of the positive GBM cases were detected, while the AUROC below 0.50 means that the model had no useful discriminative ability.

**Figure 8:**
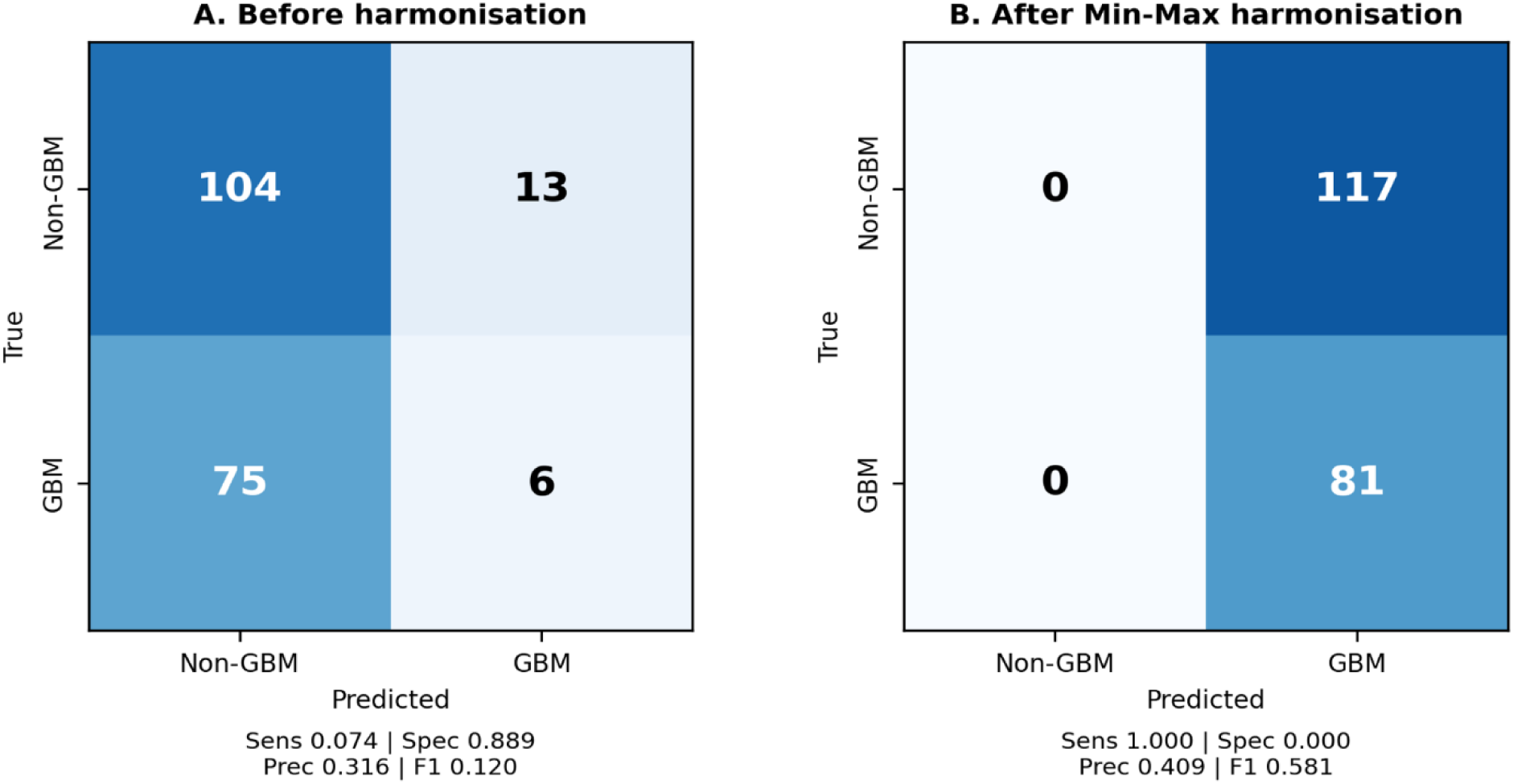
Confusion matrices of the TVAE + XGBoost champion model on CGGA (n = 1S8; 81 GBM, 117 non-GBM) (A) before and (B) after Min-Max harmonisation, annotated with sensitivity (Sens), specificity (Spec), precision (Prec) and F1-score. Class 0 denotes non-GBM glioma tissue, not healthy controls.

**Table 6:** Performance of the TVAE + XGBoost champion model on the CGGA external cohort before and after Min-Max harmonization.

| **CGGA evaluation condition** | **Accuracy** | **Recall** | **AUROC (95% CI)** |
|---|---|---|---|
| Before harmonization | 0.556 | 0.074 | 0.403 (0.324-0.482) |
| After Min-Max harmonization | 0.409 | 1.000 | 0.467 (0.386-0.548) |

The confusion matrix **(Figure 8A)** shows that only 6 of 81 GBM samples were detected, whereas 104 of 117 non-GBM samples were correctly classified (specificity 0.889). Accuracy (0.556) was below the NIR of 0.591, and the AUROC was significantly below 0.5 (0.403, 95% CI 0.324-0.482; p = 0.016), indicating that the model’s ranking of CGGA samples was systematically inverted relative to the GBM label rather than merely uninformative.

It is important to note that when the feature matrix was examined it was found that the 201 miRNA features appeared in the same order in both the training and CGGA datasets, so the failure observed could not be due to a simple mistake relating to the ordering or alignment of the feature columns.

### 3.7. Effect of Feature Harmonization on External Classification

The large difference in the ranges of the numerical features between the discovery/training data and the CGGA led to a review of Min-Max harmonization. The values in the training data were about from 0 to 47,580, while those in CGGA varied from about −129.1 to 28,912.2. Min-Max transformation was thus carried out separately on each of the datasets before the retraining and external evaluation.

After harmonization the AUROC rose from 0.403 to 0.467. Yet it was still below 0.50 and so showed no useful discriminatory ability. More significantly, harmonization greatly changed the operating characteristics of the classifier: the recall increased from 0.074 to 1.000 while the accuracy fell from 0.556 to 0.409.

The confusion matrix after harmonisation **(Figure 8B)** shows that the classifier assigned every CGGA observation (198 of 198) to the GBM class: all 81 GBM samples were detected, but none of the 117 non-GBM samples was correctly classified (specificity 0.000), so precision (0.409) and accuracy (0.409) both equalled the GBM prevalence. The rise in recall therefore reflects a shift to universal positive prediction rather than the recovery of a biologically meaningful decision boundary. Together with the fall in accuracy and an AUROC that remained below 0.50 (0.467, 95% CI 0.386-0.548), this shows that the apparent improvement in sensitivity did not correspond to improved diagnostic discrimination.

### 3.8. Generalization Across Increasingly Stringent Validation Settings

The consistent difference between improvement within the same modality and cross-domain generalization emerged from the results at the various validation levels.

The AUROCs of the non-augmented models varied between 0.421 and 0.527 when tested on the independent GSE184472 plasma cohort, while the most successful model after generative augmentation, TVAE + XGBoost, obtained an AUROC of 0.592 on the same cohort. This shows that augmentation led to a measurable but only slight improvement in discriminatory ability when validated on an independent dataset of the same modality.

The same improvement was not seen in the cross-platform CGGA cohort. When the best model was deployed directly, the AUROC obtained was 0.403, whereas Min-Max harmonization increased the value only to 0.467. Both values remained below 0.50. Moreover, the post-harmonization recall of 1.000 was accompanied by an accuracy was only 0.409, which shows that the model had tended to predict the positive class rather than having achieved better separation between GBM and non-GBM samples.

Put together, these findings show a clear gap in generalization between independent validation within the same modality and evaluation across platforms and across tissues. The small improvement seen after synthetic augmentation on GSE184472 thus did not serve as evidence of robust transferability to an externally located cohort that is both biologically and technically different.

### 3.9. Integrated Performance Across Validation Stages

The evaluation framework which was developed led to three main findings. Firstly, even though three different baseline classifiers were used, direct learning from the 38-subject discovery group gave poor external discriminative ability. Secondly, generative augmentation especially when TVAE was combined with XGBoost resulted in a small increase in AUROC on an independent plasma cohort, although the level of performance achieved was still not sufficient to support a diagnostic claim. Thirdly, this improvement did not hold up in the much more rigorous CGGA evaluation, since the AUROC values, both before and after harmonization, stayed below 0.50.

Therefore, the principal result of the study is not that CTGAN or TVAE offers a clinically effective approach to the small-cohort problem. On the contrary, the results show that the performance improvements seen after generative augmentation and assessed using the same modality during validation do not indicate the presence of a transferable diagnostic signal when the model is faced with biological and platform distribution shifts. Thus, the CGGA analysis acts as an empirical stress test demonstrating the importance of carrying out external cross-platform validation before concluding that performance gains obtained through synthetic-data augmentation amount to evidence of generalizable diagnostic ability.

## 4. Discussion

The central finding of this study is that generative data augmentation led to a small gain in glioblastoma classification when applied to an independent plasma-miRNA cohort, even though this benefit did not result in reliable cross-domain generalisation. Models that were trained directly on the 38-subject GSE122488 cohort performed badly on the independent GSE184472 plasma cohort (AUROC 0.421-0.527). Augmentation improved performance for several model combinations, with TVAE-XGBoost achieving the highest AUROC of 0.592. Although this represented an absolute improvement of 0.133 over the corresponding non-augmented XGBoost model, the performance remained insufficient for a diagnostic claim. These findings indicate that synthetic augmentation can enhance predictive performance without necessarily leading to a transferable disease signature.

The contrast between same-modality and cross-domain validation was especially informative. The TVAE-XGBoost model chosen achieved an AUROC of 0.592 on GSE184472 but just 0.403 on the independent CGGA cohort. Since CGGA differs from both the discovery and validation cohorts in terms of biological specimens and the measurement platform, there is a significant domain shift. Therefore, the decline in performance cannot be attributed to one factor alone and might be due to the combined influence of plasma-versus-tissue biology, platform-specific measurement characteristics, and cohort heterogeneity. Most importantly, the cross-domain result shows that successful validation in circulating plasma does not prove that the findings can be generalized to a different molecular measurement domain.

A further, and possibly decisive, contributor is that the classification task was not identical across cohorts. In GSE122488 the positive class comprised all tumours (12 glioblastomas and 10 grade II-III gliomas) and the negative class comprised healthy controls, whereas in CGGA the positive class was restricted to GBM (GBM, sGBM and rGBM) and the negative class consisted of all other glioma histologies. A model that learned a general glioma-versus-healthy signal from plasma would therefore be expected to score many lower-grade CGGA tumours as positive. CGGA consequently tests the discrimination of GBM from other gliomas, which is a harder and clinically different question from the detection of glioma against healthy controls. The near-universal positive predictions after harmonisation and the AUROC significantly below 0.5 before harmonisation (p = 0.016) are both consistent with this label-definition mismatch. Future cross-cohort evaluations should align the label definition across cohorts, for example by restricting the discovery positive class to GBM, because CGGA contains no non-tumour tissue against which a glioma-versus-healthy rule could be tested.

The biological compartment also differs. Circulating plasma miRNAs originate from many tissues, including blood cells, and their measured levels are influenced by haemolysis and pre-analytical handling, whereas tissue profiles reflect the intracellular miRNA content of the tumour and its microenvironment. Concordance between tissue and circulating levels of individual miRNAs is therefore not guaranteed, and a plasma-derived decision rule has no a priori reason to transfer to tissue even when the same miRNAs are measured. Platform differences add a technical shift on top of this biological one: the training data were non-negative (0 to 47,580), whereas CGGA contained negative values (minimum −129.1), indicating a different normalisation or background-correction pipeline that alters both the scale and the shape of the expression distributions, as well as the detection limits for low-abundance miRNAs.

Per-cohort Min-Max scaling addresses only part of this shift. It aligns the minimum and maximum of each feature but not the central tendency or shape of its distribution, it is sensitive to outliers, and because it is fitted within each cohort it maps each cohort’s own extremes to 0 and 1 regardless of differences in class composition. More suitable candidates for future work include rank-based or quantile normalisation, within-sample features such as miRNA expression ratios that are invariant to global scale, and recalibration of the decision threshold on a small labelled subset of the target domain. Batch-correction methods such as ComBat would be confounded in this setting because the plasma and tissue cohorts do not share comparable control groups.

Finally, the size of the evaluation cohorts limits statistical resolution. With 45 tumour and 73 control samples, the 95% CI around an AUROC of about 0.6 spans roughly ±0.11, and none of the AUROCs observed on GSE184472 could be distinguished from chance; the 0.133 gain from TVAE augmentation did not reach significance under a conservative comparison (p = 0.084). All operating points were also evaluated at a fixed 0.5 threshold, whereas the tumour-enriched synthetic class prior of TVAE (75.8% tumour) would be expected to shift predicted probabilities upwards; this plausibly contributes to the tendency towards positive predictions in both external cohorts. Threshold selection or probability recalibration on independent data would be required before any operating point could be interpreted clinically.

The fact that the synthetic-data distributions have been taken into account should be noted. The dataset produced by CTGAN was fairly balanced (253 tumor cases and 247 control cases), while that generated by TVAE was considerably enriched in tumor cases (379 tumor cases and 121 control cases). This imbalance might have affected the decision boundary that was learned and could therefore in part account for the strong positive-class predictions seen in the external testing. Nevertheless, the current analysis does not prove a causal link between the imbalance in the synthetic classes and the cross-domain failure. Instead, it underscores the importance of actively controlling and assessing the distributions of the synthetic classes when augmentation is carried out on small biomedical cohorts.

Even though feature scaling alone was insufficient to overcome external discrimination, min-max harmonization raised the CGGA performance from an AUROC of 0.403 to 0.467 and increased recall from 0.074 to 1.000, while accuracy fell from 0.556 to 0.409. The higher recall thus indicated a shift towards making mostly positive predictions rather than an improvement in the separation of the classes. This result implies that the numerical differences in scale were not the main obstacle to transfer in the situation under investigation; after simple scaling, distributional and biological differences probably still remained. More thorough cross-domain harmonization and validation will be needed in order to find out whether transferable molecular signals can be recovered.

The feature ranking derived from the model indicated that hsa-miR-301b was the most important feature (with a Gini importance of 0.183), after which came hsa-miR-760 (0.049). However, these results are hypothesis-generating and not proof that these are validated biomarkers, especially since the discovery cohort was small and had limited ability for external discrimination. Only when the features have been shown to be stable across resampling, in independent cohorts, and through biological validation can they be given diagnostic significance.

### Limitations

The discovery cohort was small and heterogeneous (GBM and lower-grade gliomas pooled as tumour); each generator and classifier was trained with a single random seed, so run-to-run variability was not quantified; generator hyperparameters were not tuned; class labels were not defined identically across cohorts; GSE90603 was not used for an additional external test; and predicted probabilities were not calibrated. These limitations reinforce the conclusion that the present results do not support a diagnostic claim.

## 5. Conclusion

The study shows that generative data augmentation can lead to better classification of GBM in a small cohort of circulating-miRNAs, but the improvement seen does not mean that robust external generalisation will result. For the independent plasma cohort of the same modality, augmentation increased performance, the TVAE-XGBoost model obtaining an AUROC of 0.592 as compared to the range of 0.42 to 0.53 for the models that had not been augmented. Yet, performance dropped significantly in the independent CGGA cohort, the AUROC being 0.403 before and 0.467 after Min-Max harmonisation. It therefore shows that the performance improvements seen in the small discovery group in this study cannot be taken as proof of the diagnostic generalizability. The study also proves the need to assess molecular classifiers in ways that go beyond validation within the same modality before one can consider their potential for translation or diagnostic use. The miRNA features identified, such as hsa-miR-301b, still remain as potential signals and thus require further independent validation. In summary, this research offers a practical foundation for the rigorous external evaluation of generative-augmentation methods in small-cohort biomedical machine learning.

## Authors’ Contribution

PKM devised the concept, developed the methodology, and supervised the in silico experiments; SR, SM, and PC performed most of the in silico experiments; SM, PC, and VG performed the data analysis and interpretation; RT reviewed the manuscript; and SR, SM, VG, and PKM drafted the original manuscript.

## Data Availability

The data supporting this study’s findings are available upon request from the corresponding author.

## Abbreviations

GBM: Glioblastoma multiforme
TVAE: Tabular Variational Autoencoders
CTGAN: Conditional Tabular Generative Adversarial Network
AUROC: Area Under the Receiver Operating Characteristic curve
CGGA: Chinese Glioma Genome Atlas

